# Levamisole suppresses activation and proliferation of human T cells by the induction of a p53-dependent DNA damage response

**DOI:** 10.1101/2023.05.08.539819

**Authors:** Gerarda H. Khan, Floor Veltkamp, Mirte Scheper, Ron A. Hoebe, Nike Claessen, Loes Butter, Antonia H.M. Bouts, Sandrine Florquin, Jeroen E.J. Guikema, the LEARNS Consortium

## Abstract

Levamisole (LMS) is a small molecule used in the treatment of idiopathic nephrotic syndrome (INS). The pathogenesis of INS remains unknown, but most evidence points towards an immunological basis of the disease. Recently, LMS has been shown to increase the relapse-free survival in INS patients treated in combination with corticosteroids with relatively few side effects. While LMS has been hypothesized to exert an immunomodulatory effect, its mechanism of action remains unknown. To provide insight into the working mechanism of LMS, we studied its immunomodulatory activity on in vitro activated human T cells. We show here that treatment with LMS decreased activation and proliferation of human CD4+ and CD8+ T cells. In addition, production of T cell activation-associated cytokines such as IL-2, TNF-α and IFN-γ were reduced upon LMS treatment, whereas IL-4 and IL-13 production was increased. Gene expression profiling confirmed the suppressive effects of LMS on proliferation as numerous genes involved in cell cycle progression were downregulated. Furthermore, genes associated with p53 activation and cell cycle arrest were upregulated by LMS. In agreement, LMS treatment resulted in p53 phosphorylation and increased expression of the p53 target gene FAS. Accordingly, LMS sensitized activated T cells for Fas-mediated apoptosis. Cell cycle analysis showed that LMS induced a mid-S phase arrest indicating the activation of a replication stress-associated checkpoint. In support, LMS treatment resulted in γH2AX-foci formation and phosphorylation of CHK1. Our findings indicate that LMS acts as an immunosuppressive drug that directly affects the activation and proliferation of human T cells by induction of DNA damage and the activation of a p53-dependent DNA damage response.

## 1. INTRODUCTION

Levamisole (LMS) is a synthetic imidazole derivate and small molecule that has been described to have a wide range of immunomodulative properties (1). LMS is a known inhibitor of tissue non-specific alkaline phosphatase (TNAP) and an agonist of the nicotinic acetylcholine receptor (nAChR) (2, 3), but whether these activities underlie its immunomodulatory action remain unknown. Clinical activity of LMS in the treatment of multiple immune-mediated diseases supports its immunomodulatory properties (4, 5). Recently, LMS has been shown to reduce the relapse rate in relapsing idiopathic nephrotic syndrome (INS) (6-10), a kidney disease characterized by proteinuria, edema and hypoalbuminia (11).

T cells play a prominent role in the majority of immune-mediated diseases (12, 13), and are the prime therapeutic targets of established drugs that are used for the treatment of autoimmune diseases, such as mycophenolate mofetil, methotrexate and calcineurin inhibitors (14-18). INS has been recognized as a T cell-mediated disorder, although its etiology remains unknown (19-21). The LEARNS (LEvamisole as Adjuvant therapy to Reduce relapses of Nephrotic Syndrome) consortium, of which our research group is part of, is currently conducting a randomized control clinical trial assessing the effects of LMS on the relapse rate and immune-related parameters in newly diagnosed pediatric INS patients (22).

The mechanism by which LMS affects immune cells remains largely unknown. Several studies have reported on the effects of LMS on T cells, and suggested that LMS promotes a T-helper 2 cell (Th2) to Th1 shift, primarily based on altered cytokine production profiles (23-26). In addition, it was shown that LMS may either increase T cell proliferation (27-29) or diminish it (27, 30), depending on the context, drug concentration, and species (human versus mouse) that are studied. More recently, it was shown that LMS decreased *in vitro* proliferation of human bone marrow cells and CD4+ T cells, which was hypothesized to occur through its effects on Toll-like receptor (TLR) and JAK/STAT signaling (31). However, whether reduced JAK/STAT signaling in LMS treatment initiates the observed T cell suppression remains unclear.

As of yet, a comprehensive and in-depth analysis of the exact mode of action of LMS on activated human T cells is lacking. The aim of this study is to provide further insight into the effects of LMS on T cell activation and proliferation. Here, we demonstrate that LMS directly suppresses the activation and proliferation of activated human CD4+ and CD8+ T cells. We show that LMS suppressed T cell activation by the induction of replication stress, which triggered the activation of p53. In addition, we show that LMS sensitized T cells for the extrinsic apoptosis pathway, involving the p53-dependent induction of Fas. Our results suggest that LMS acts as an immunomodulatory drug by triggering the p53-dependent suppression of T-cell activation and proliferation, underlining its therapeutic use as an immunosuppressant drug.

## 2. MATERIALS AND METHODS

### 2.1. Isolation and in vitro culturing of primary human T cells

Human peripheral mononuclear blood cells (PBMC) were isolated from 6 buffy coats by Ficoll-Hypaque density gradient centrifugation. All buffy coats were collected from adult anonymous healthy blood donors (Sanquin Blood Supply, Amsterdam, The Netherlands) who provided written informed consent for usage of the remainder of their blood donation in research. Untouched T cells were isolated from PBMCs by negative Magnetic-activated cell sorting (MACS) using the human pan T cell isolation kit (Miltenyi Biotec, Bergisch Gladbach, Germany) according to the manufacturer’s instructions. Purity of the T cell fraction was assessed by flowcytometry for CD3 (>98% for all donors). Purified T cells were cultured in Roswell Park Memorial Institute (RPMI) medium supplemented with 2mM L-glutamin, 100 U/mL of penicillin, 100 μg/mL of streptomycin (Gibco, Thermo Fisher Scientific, Waltham, MA, USA), and 10% fetal calf serum (FCS; HyClone, GE Healthcare Life Sciences, Chicago, IL, USA). T cells were cultured at a density of 1×10^6^ cells/mL at 37 °C/5% CO2. Purified T cells were activated for 1 to 3 days with Human T-Activator anti-CD3/CD28 dynabeads at a 1:2 bead-to-cell ratio (Gibco, Thermo Fisher Scientific, Waltham, MA, USA). Prior to analyses, the T cells were detached from the dynabeads by manual dispersion and magnetic separation, as per the manufacturer’s instructions. At the start of activation, T cell samples were treated with 1mM levamisole (levamisole-hydrochloride, Sigma-Aldrich, St. Louis, MO, USA), 1 μM doxorubicin (Doxorubicin hydrochloride, Sigma-Aldrich) or 1:200 μL H_2_O (solvent control condition). Levamisole was added at the start of the culture and refreshed after 24 hours. In indicated experiments, samples were also treated with 10 μM Quinoline-Val-Asp-Difluorophenoxymethylketone (Q-VD-Oph, Selleckchem, Houston, TX, USA) and/or 10 μM hexameric FAS receptor agonist APO010 (a kind gift from Prof. dr. Eric Eldering, Amsterdam UMC).

### 2.2. Gene expression profiling by RNA sequencing

RNA sequencing was performed on activated human peripheral blood T cells obtained from 3 healthy donors. Total T cells (CD3+) were activated for 24 hours and 48 hours with anti-CD3/CD28 and treated with 1 mM LMS or solvent (untreated controls). RNA was isolated using TRIzol (Thermo Fisher Scientific) and the RNeasy Mini Kit (Qiagen, Hilden, Germany) according to the manufacturer’s instructions. RNA quality was assessed by determining the RNA Integrity Number (RIN) on a TapeStation system (agilent Technologies, Santa Clara, CA, USA). All RNA samples had a RIN higher than 7, with a mean RIN of 9.76. Sequencing libraries were constructed by using the TruSeq stranded mRNA library kit and paired-end sequenced by the Illumina HiSeq platform. Quality control, trimming, alignment and quantification were all performed using the Galaxy web platform (https://usegalaxy.org/) (32). All reads had a PHRED score >28 (mean PHRED score of 36). The resulting normalized feature Counts table was both uploaded to and analyzed by using R2: Genomics Analysis and Visualization Platform (http://r2.amc.nl) and analyzed in R studio. Differential expression was analyzed using DESeq2 (33). Differentially expressed genes were defined as genes with an absolute log2 fold change (Log2FC) greater than 1 with a p-value of <0.05. Enrichment plots were created by using the Broad institute Gene Set Enrichment Analysis (GSEA) software (34).

### 2.3. Flow cytometric analyses

Activated T cells were washed with FACS buffer (0.2% BSA, 0.5 mM EDTA, 0.1% sodium azide in Phosphate buffered saline, PBS) and stained with the respective antibodies/dyes for 30 minutes on ice while protected from light. All measurements were performed on a BD LSRFortessa and data was analyzed using FlowJo™ (v10.5 Software, FlowJo LLC, Ashland, OR, USA). Gates were drawn based on Fluorescence minus one (FMO) stainings. The following antibodies/dyes were used: CD3-APC (SK7, Becton Dickinson,Franklin Lakes, NJ, USA), CD4-PB (RPA-T4, Biolegend,San Diego, CA, USA), CD8-APC (SK1, Biolegend), CD69-PE (L78, Becton Dickinson), 7-aminoactinomycin-D (7-AAD; BioLegend), FAS-PEcy7 (DX2, CD95, Biolegend), CellTrace carboxyfluorescein succinimidyl ester (CFSE) (Invitrogen, Thermo Fisher Scientific), CaspGLOW Fluorescein Active Caspase-3 Staining Kit (Thermo Fisher Scientific), Phospho-p53 (Ser15)-PE (16G8, Cell signaling Technology, Danvers, MA, USA). Phosphorylation of p53 was measured by fixation of the cells with 4% paraformaldehyde (PFA), followed by permeabilization with ice-cold 90% methanol and staining with pP53-PE for 1 hour at room temperature. Cell cycle analysis was performed by using the bromodeoxyuridine (BrdU) incorporation method as described previously (35). In short, cells were incubated for 1 hour with 40 μM BrdU (Thermo Fisher Scientific) after which the cells were fixed in 75% cold ethanol. After fixation the cells were digested using pepsin and HCl. BrdU incorporation was assessed by staining with anti-BrdU-FITC (Becton Dickinson, #347583) and TO-PRO™-3 (Thermo Fisher Scientific) and analyzed by flow-cytometry.

### 2.4. γ-H2AX foci analysis

For the analysis of γ-H2AX foci, 48 hours activated T cells were adhered to poly-L-Lysine-coated coverslips. 12 mm round coverslips were sterilized and coated with 10 μg/mL of Poly-L-Lysin (Poly-L-Lysine Solution (0.01%), Sigma-Aldrich) and washed with PBS. 50 μL of T cell suspension in PBS was pipetted on the coverslips and left for 30 minutes at room temperature to attach. After attachment, the T cells were fixed with 4% PFA, blocked and permeabilized with PBT buffer (0.1% TritonX-100, 0.5% BSA in PBS). γ-H2AX staining was performed in PBT buffer in a 1:10,000 dilution (gamma H2AX Ser139, Rabbit, Novus Biologicals, Englewood, CO, USA) and incubated overnight. The coverslips were then stained with Alexa fluor 594 donkey-anti rabbit 1:250 (Donkey anti-Rabbit IgG ReadyProbes™, Alexa Fluor™ 594, Invitrogen, Thermo Fisher Scientific) for 2 hours. Nuclei were stained with Hoechst33342 1:2000 (Hoechst 33342 Solution (20 mM), Invitrogen, Thermo Fisher Scientific) for 1 hour after which the coverslips were mounted on slides with aqua-poly/mount (Aqua-Poly/Mount, Polysciences, Warrington, PA, USA) and analyzed by fluorescence microscopy. Z-stack images were collected on the Leica Thunder Wide Field Fluorescence Microscope and were deconcolved by using Huygens deconvolution software (Scientific Volume Imaging, The Netherlands, http://svi.nl). Maximum intensity projections were made and were used to detect nuclei and γ-H2AX foci. Cellpose (36) was used to distinguish nuclei, while an algorithm based on StarDist (37) and trained to recognize γ-H2AX foci in primary T cells was used to distinguish γ-H2AX foci. The percentage of cells possessing >10 nuclear foci was calculated by analyzing 6 fields of view per sample, containing at least 350 cells per sample.

### 2.5. Immunoblotting

Immunoblotting was performed as previously described (38). The following antibodies were used: Phospho-p53 (Ser15) (Rabbit, Cell signaling Technology, #9284), p53 (7F5) (Rabbit, Cell signaling Technology, #2527), Phospho-Chk1 (Ser317) (Rabbit, Cell signaling Technology, #12302), Chk1 (2G1D5) (Mouse, Cell signaling Technology, #2360), β-Actin (8H10D10)(Mouse, Cell signaling, #3700).

### 2.6. Cytokine multiplex assay

Concentrations of cytokines in supernatants of T cell cultures were assessed with the MILLIPLEX MAP Human Th17 magnetic bead panel kit (Merck Millipore, Burlington, MA, USA) and the Luminex® 200™ system according to the guidelines of the manufacturer. In short, supernatants of 72 hours activated T cells samples from 6 T-cell donors were collected and stored at -80°C. After thawing, samples were centrifuged at 1000xg for 10 minutes. Cytokine concentrations were calculated based on the median fluorescent intensity (MFI) of the sample compared to the MFI of the standard curve.

### 2.7. Statistics

Statistical analysis was performed by using GraphPad prism software (Graphpad Software, version 8.0.0 for windows). Median fluorescence indexes and geometric means of fluorescence were calculated with FlowJo™ Software (version 10). Division indexes were also calculated in FlowJo by using the proliferation analysis module on CFSE stainings.

## 3. RESULTS

### 3.1. Levamisole inhibits proliferation and activation of human peripheral blood CD4+ and CD8+ T cells

To assess the effect of LMS treatment on T cell proliferation *in vitro*, total human peripheral blood T cells were stained with CFSE and activated with anti-CD3/CD28. Proliferation was significantly reduced by treatment with 1 mM LMS for 72 hours (Fig 1A), but remained unaffected by lower concentrations (1 μM – 100 μM) of LMS (Supplemental fig 1C). Flow cytometry analysis showed that LMS inhibited proliferation of cytotoxic T cells (CD8+) and T helper cells (CD4+) to a comparable degree (Fig 1A, Supplemental Fig 1D). Importantly, live/dead flow cytometry staining showed that LMS treatment did not induce cell death of dividing T cells (Supplemental Fig 1C). In addition to its inhibitory effects on proliferation, LMS treatment resulted in a significantly lower expression of the early activation marker CD69 (39) in both CD4+ and CD8+ T cells (Fig 1B and 1C). In support, the secretion of pro-inflammatory cytokines that are associated with T-cell activation, such as IL-2, IFNγ and TNFα, were significantly reduced upon LMS treatment. Suprisingly, IL10 was also significantly reduced, whereas IL4 and IL13 were significantly increased in the supernatants of LMS-treated T cells (Fig 1D). These results indicate that LMS affects T cells directly and suppresses T-cell proliferation while simultaneously stimulating the production of Th2 associated cytokines.

**FIGURE 1:**
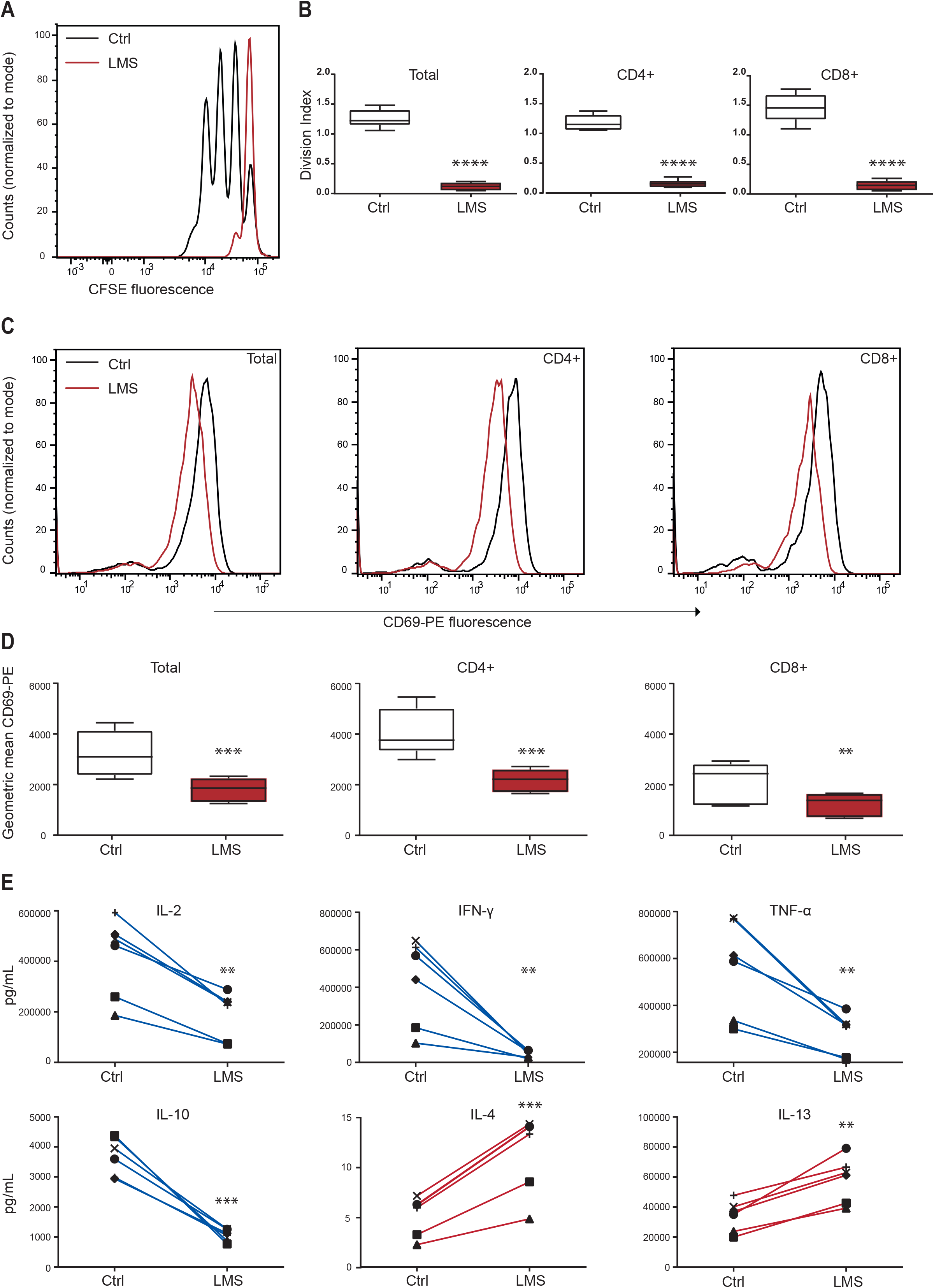
Decreased proliferation, activation and cytokine production in LMS-treated human T cells. **(A)** LMS treatment decreased proliferation of anti-CD3/CD28 bead-activated T cells, as measured by the CFSE proliferation assay. T cells were stained with CFSE and activated with anti-CD3/CD28 dynabeads in a 2:1 cell:beads ratio for 72 hours and treated with 1 mM LMS or solvent. **(B)** Division indices were calculated for (from left to right) total T cells (CD3+), CD4+, and CD8+ T cells (n = 6 donors). Mean and SD of the division indices of 6 different donors are shown (***P< .0001; paired t test). **(C)** LMS treatment decreased the expression of the early T cell activation marker CD69. Flow cytometry histograms of CD69 expression for solvent-treated (black) and LMS-treated T cells (red), activated for 24 hours are shown. (**D)** Graphs depicting fluorescence geometric means of CD69 staining in solvent treated activated T cells (open boxes) and LMS-treated activated T cells (red boxes). Mean and SD are shown (n = 6 donors) (**P < .001, ***P < .0001; paired t test). **(E)** LMS treatment decreased the production of IL-2, IFN-γ, TNF-α and IL-10, and increased the production of IL-4 and IL-13. Supernatants were harvested from anti-CD3/CD28 stimulated T cell cultures (72 hours stimulation). Cytokine concentrations were measured by the Milliplex™ assay. The calculated concentration in pg/mL of the individual donors is shown here (n = 6 donors) (**P < .001, ***P < .0001; paired t test).

### 3.2. Levamisole affects the transcription of genes involved in cell-cycle progression and DNA replication

To investigate the potential mechanisms and targets of LMS we analyzed its effects on gene expression in human T cells from three individual healthy donors by RNA sequencing. Importantly, principal component analysis (PCA) of gene expression profiles obtained after 24 hours and 48 hours of treatment indicated that LMS treatment is a major determinant of variation in gene expression, showing the most prominent differential expression at 48 hours (Supplemental Fig 2A and B). Using a Log2FC >1 and a p-value <0.05 as cutoff, 346 differentially expressed genes were identified. Of these differentially expressed genes, 123 genes were downregulated and 223 genes upregulated in LMS treated T cells. In agreement with its effect on T-cell proliferation, numerous genes involved in cell-cycle progression and T cell expansion such as *CCNB1, PLK1* and *IL2* were found to be downregulated (Fig 2A). Correspondingly, gene set enrichment analyses (GSEA) confirmed the enrichment of genes involved in cell-cycle and DNA replication in untreated activated T cells over LMS-treated T cells (Fig 2B). Similarily, several genes associated with cell cycle arrest and p53 activation such as *TP53INP1* and *CDKN1A* were upregulated in LMS-treated T cells (Fig 2A). Moreover, GSEA showed the significant enrichment for gene sets involved in p53 activation in LMS-treated T cells (Fig 2B).

**FIGURE 2:**
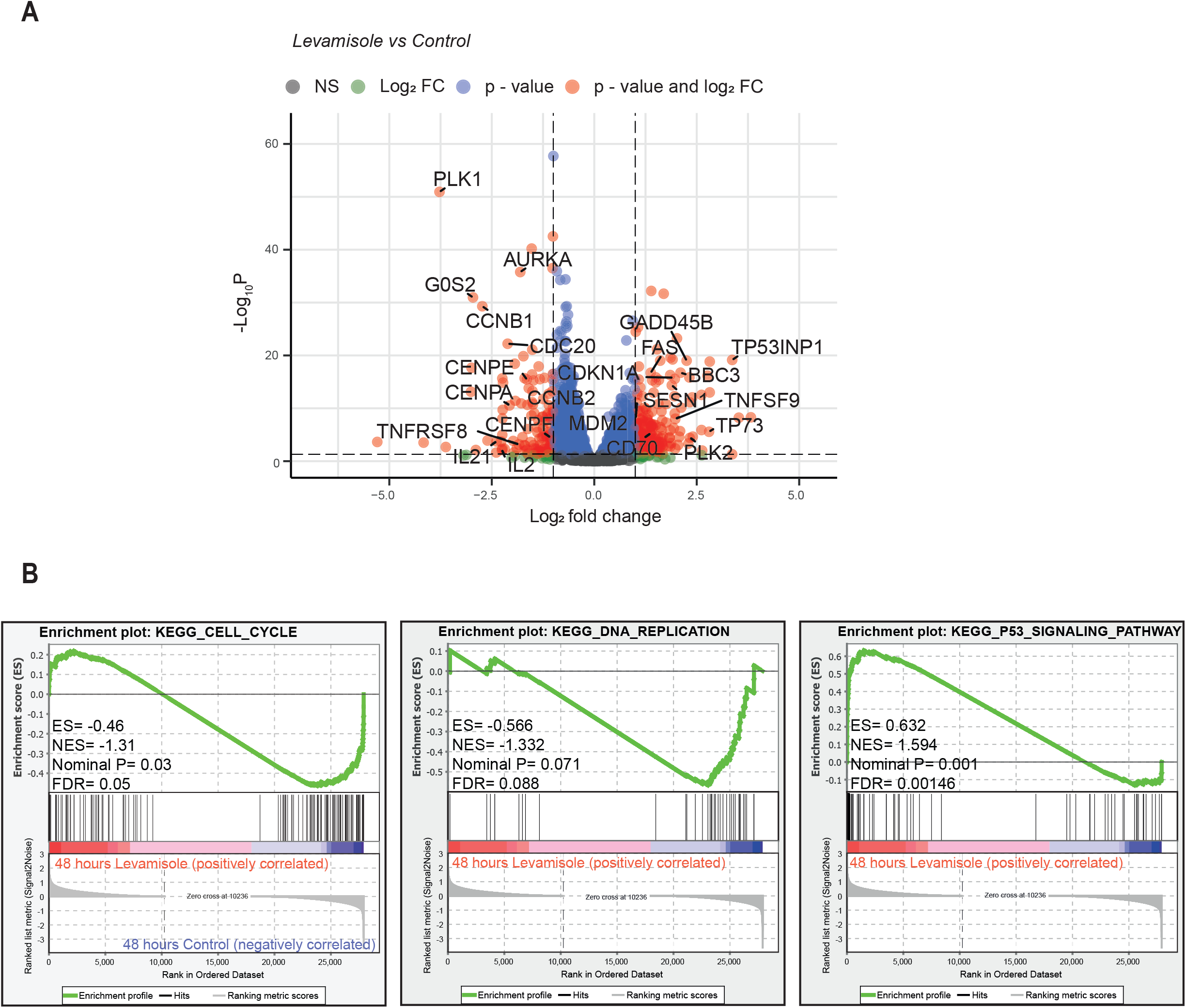
Gene expression profiling of LMS-treated activated human T cells shows downregulation of cell cycle-associated genes and upregulation of p53 target genes. **(A-B)** Human peripheral blood T cells (CD3+) from 3 donors were activated for 48 hours with anti-CD3/CD28 dynabeads and treated with LMS 1 mM or solvent. Gene expression was assessed by RNA sequencing. Differentially expressed genes were defined as genes with an absolute Log2FC greater than 1 and with a p-value of <.05. (A) Volcano plot showing differentially expressed genes comparing LMS-treated samples to solvent-treated samples. Genes involved in cell cycle progression and p53 activation are indicated. **(B)** GSEA enrichment plots show enrichment of genes involved in cell cycle progression in solvent-treated T cells over LMS-treated T cells. GSEA enrichment plots for (from left to right) KEGG_CELL_CYCLE, KEGG_DNA_REPLICATION and KEGG_P53-SIGNALING_PATHWAY. False discovery rate (FDR), enrichment score (ES), normalized enrichment score and p-value are shown in the individual plots.

### 3.3. Levamisole treatment results in p53 activation and sensitization for the extrinsic apoptosis pathway in activated human T cells

In further support of the effect of LMS treatment on the activation of p53 in T cells, we demonstrate by immunoblotting that LMS resulted in the phosphorylation of p53 at residue serine 15 (Ser15), which provokes p53 stabilization and is crucial for the induction of cell-cycle arrest (40)(Fig 3A-C). Moreover, we show that the expression of the p53 target gene *FAS*, which codes for the Fas receptor that is involved in the extrinsic apoptosis pathway, was significantly increased in LMS treated T cells (Fig 3D,E). In addition, LMS treated T cells showed a higher expression of Fas (Fig 3F) in both CD4+ and CD8+ T cells as shown by flow cytometry (Fig 3E), whereas the expression of the Fas ligand (Fas-L, CD95L) was unchanged (Supplemental Fig 3A and B). To assess whether the increased expression of Fas sensitized T cells for the induction of the extrinsic apoptosis pathway, cells were treated with a synthetic Fas ligand analogue (FAS10) and active (cleaved) caspase-3 was measured using the CaspGLOW assay. FAS10 induced caspase-3 cleavage in both the control and LMS treated samples but this increase was significantly higher in the LMS treated samples, suggesting that LMS treatment sensitizes T cells for Fas-induced apoptosis (Fig 3G). Furthermore, treatment with the pan-caspase inhibitor Q-VD-Oph prevented caspase-3 cleavage and FAS10-induced apoptosis, whereas it had no effect on the inhibition of proliferation by LMS (Fig 3H).

**FIGURE 3:**
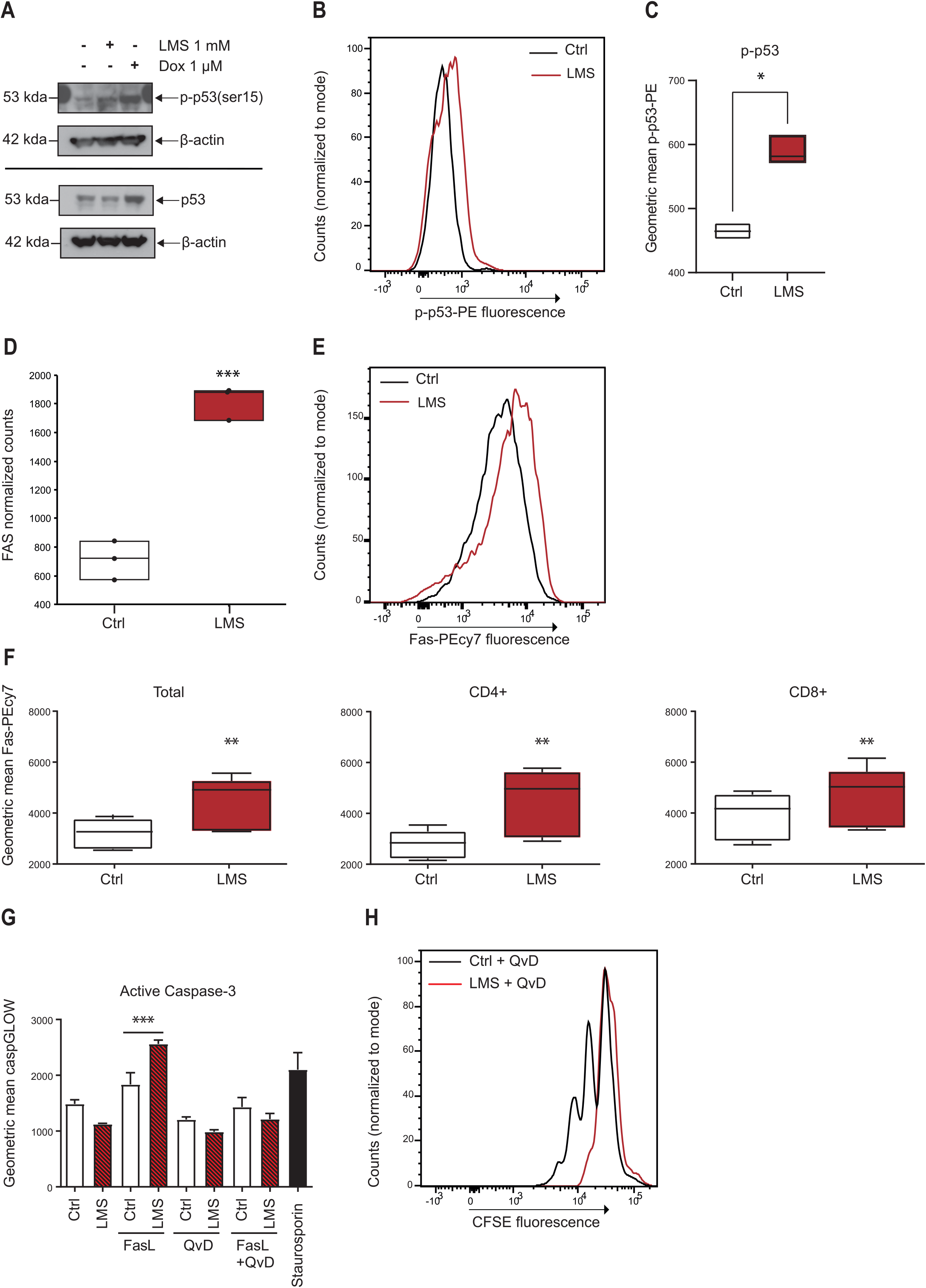
LMS induces the activation of p53 and sensitizes T cells for FAS-mediated apoptosis. **(A)** Immunoblot analysis of phosphorylated p53 (ser15) and total p53 protein levels in control-, LMS- and doxorubicin-treated T cells after 48 hours of activation (n = 1 donor). β-actin was used as a loading control. **(B)** Intracellular flow cytometry staining for phosphorylated p53 (Ser15) in solvent-treated T cells (Ctrl, black histogram) and LMS-treated T cells (red histograms). T cells were activated and treated with LMS for 48 hours **(C)**. Graphs depicting the geometric means of fluorescence for the phospho-p53 staining (n = 3 donors) (*P < .01; paired t test). **(D)** Graph showing the normalised RNA-seq counts for the p53 target gene FAS (n =3 donors) (*** P<.0001; One Way Analysis of variance, ANOVA). **(E)** Flow cytometry staining for Fas receptor (Fas, CD95) on activated human T cells treated with 1mM LMS (red histogram) or solvent (Ctrl, blac histogram) for 72 hours **(F)** Graphs depicting the fluorescence geometric means Fas staining in total (CD3+), CD4+ and CD8+ T cells (n = 6 donors) (**P<.001; paired t test). **(G)** Graph depicting the geometric means of fluorescence for the active caspase-3 staining (caspGLOW). T cells were treated with 1 mM LMS or solvent controls for 48 hours after which the medium was changed and the cells were treated for 48 hours with either 5 μM FAS10 (a Fas-ligand), 10 μM QvD-OPH (pan-caspase inhibitor), or with the combination of both. As a positive control 10 μM Staurosporin was used (n=3 donors, ***P<.0001; One Way Analysis of variance with Bonferroni’s multiple comparison test). **(H)** Flowcytometry histogram of CFSE fluorescence for activated T cells treated with 10 μM QvD-OPH (black) and activated T cells treated with both 1 mM LMS and 10 μM QvD-OPH for 72 hours (red). Treatment with the pan caspase inhibitor QvD-OPH did not affect the inhibition in cell division observed in LMS-treated T cells (n = 1 donor)

### 3.4. Levamisole causes an intra S-phase cell-cycle arrest by inducing DNA damage

To further investigate the effects of LMS on the cell-cycle progression in activated T cells, BrdU incorporation assays were performed. Our results indicate that LMS induced a cell-cycle arrest at the mid-S phase, suggesting that LMS prevented cell-cycle progression during DNA synthesis by provoking replication stress, resulting in the activation of an intra-S-phase checkpoint. Additionally, an accumulation of cells in G1 was observed, which may indicate that cells with DNA damage that proceed to undergo mitosis subsequently fail to pass the G1- to S-phase checkpoint (Fig 4A and B). In agreement, the phosphorylation of the checkpoint kinase 1 (Chk1) at serine 317 (Ser317), which is critically involved in cell-cycle checkpoints and the cellular response to DNA damage (41), was increased in activated T cells upon LMS treatment (Fig 4C). To examine whether LMS induced DNA damage in activated T cells, the formation of γ-H2AX foci was assessed. Phosphorylation of the serine 139 (Ser139) residue of the histone variant H2AX, forming γ-H2AX, is an early cellular response to in the induction of DNA double strand breaks (DSB), which is involved in the recruitment of repair proteins (42, 43). LMS-treated activated T cells showed significantly more nuclear γ-H2AX foci compared to untreated activated T cells (Fig 4D and E). These findings suggest that LMS induced DNA damage in activated T cells, resulting in a cell-cycle arrest.

**FIGURE 4:**
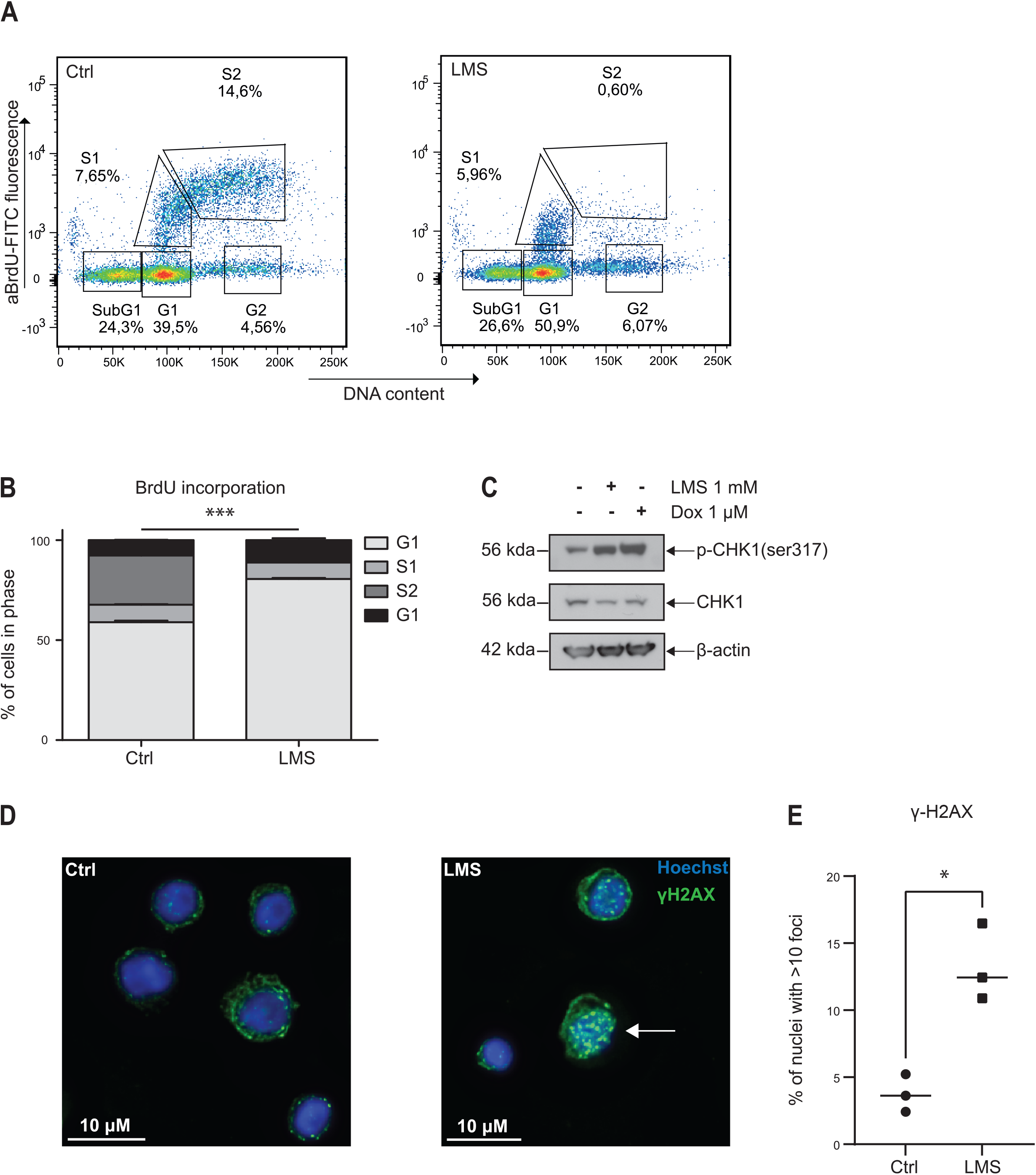
LMS causes replication stress by induction of DNA damage. **(A)** Bromodeoxyuridine (BrdU) incorporation cell cycle analysis depicting the cell cycle phases: sub-G1 (dead cells), G1, S1 (early S phase), S2 (late S phase) and G2. Activated T cells were treated with 1 mM LMS or solvent (Ctrl) for 48 hours. Cells were analysed for BrdU and DNA content by flowcytometry. **(B)** Bar graph depicting the percentages of cells in the different cell cycle phases. Sub-G1 phase cells were excluded from the analysis and percentages were calculated for the remaining cell cycle phases (n = 3 donors) (columns were compared across the treatment groups: G1 and S2 ****P<.00001, G2 * P<.01; Two Way Analysis of variance with Šidáks multiple comparison test). **(C)** Immunoblot analysis showing the protein expression of phosphorylated CHK1 (Ser 317) and total CHK1 in control, LMS- and doxorubicin-treated T cells after 48 hours of activation (n = 1 donor). 1 μM Doxorubicin was used as a positive control. β-actin was used as a loading control. **(D)** Immunofluorescence staining of phosphorylated H2AX (green) (γ-H2AX, ser319) in human activated T cells treated with solvent (Ctrl) or 1 mM LMS for 48 hours. Nuclei were counterstained with Hoechst 33342 (blue). **(E)** Graph depicts the percentage of cells with >10 nuclear γ-H2AX foci (n = 3 donors, 350 cells assessed)(* P<.01; paired t test).

## 4. DISCUSSION

The exact nature of the immunosuppressant activity of LMS in the treatment of immune-related diseases, including INS, has remained elusive until thus far. In this study we provide evidence that LMS directly suppresses T-cell activation and proliferation by the induction DNA damage, triggering a DNA damage response that involves the activation of p53 and cell-cycle arrest. Our results suggest that the level of DNA damage induced by LMS is modest, as extensive DNA damage typically triggers apoptosis, which we did not observe. However, we show that LMS primed activated T cells for Fas-mediated apoptosis, which could contribute to the immunesuppressive activity of LMS. Importantly, LMS did no differentially affect CD4+ versus CD8+ T cells, in line with its general effect on the DNA damage response. Interestingly, LMS has previously been used in combination with the cytotoxic chemotherapeutic drug 5-fluorouracil (5-FU) for the treatment of colorectal cancer, where it was most effective in patients with unaffected p53 signaling (44, 45). The antitumor activity of 5-FU is partly attributable to its ability to induce p53-dependent cell-cycle arrest and apoptosis. LMS combined with 5-FU was shown to reduce the relapse rate and improve overall survival of resected stage III colon cancer (46, 47). Based on our results, we speculate that LMS and 5-FU exert an antitumor effect by cooperative activation of p53, suggesting that patients without p53 alterations would benefit most from this combination.

Multiple potent and widely used immunosuppressive agents such as cyclophosphamide, methotrexate and mycophenolic mofetil are known DNA damage inducers (48, 49). Other more T cell specific immunosuppressants such as cyclosporine and tacrolimus have also been associated with increased DNA damage *in vitro* (50). These immunosuppresive drugs are all associated with supressing the production of the activation associated cytokines, IL-2, IFNγ and TNFα (51-54), which is in accordance with our findings. We show that LMS greatly reduced expression of most activation-induced cytokines, in further support of the suppressive effects of LMS on T cell activation. In contrast, IL-4 and IL-13 production were found to be significantly increased in LMS treated T cells. This was surprising because LMS has previously been associated with a reduction of IL-4 production in T cells (23, 24, 26). In T cells, IL-4 and IL-13 are both produced by Th2 cells and are key cytokines in the type II inflammatory pathway. IL-4 supports B cell differentiation and promotes the differentiation of naïve CD4+ T cells into Th2 cells. IL-13 promotes IgE class switching (55). An increase of Th2-associated cytokines may point towards Th2 skewing, which typically occurs through the increase of *GATA3*, the genetic lineage marker for Th2 differentiation. However, in our data set we observed no significant increase in Th lineage markers (data not shown) making a LMS-mediated skewing towards Th2 less likely. A possible explanation for the increase in IL-4 and IL-13 in our experiments is that both are associated with p53 signaling. IL-13 has been shown to reduce p53 signaling and expression of caspase-3 in activated T cells (56) while IL-4 has been associated with both suppression and activation of p53 signaling(57, 58).

In previous studies LMS has been shown to dampen proliferation in T cells. Hernández-Chirlaque et al. (2017) showed that LMS reduced proliferation in murine T cells both *in vitro* and *in vivo* through its function as a TNAP inhibitor (30). While murine T cells express *ALPL*, the gene encoding for TNAP, human T cells do not (59). In agreement, we did not detect *ALPL* expression by RNA-seq, indicating that the measured effects are TNAP independent. More recently, Wang et al. (2022) showed that LMS reduces proliferation of human CD4+ T cells (31). The authors showed in whole bone marrow mononuclear cells and in human and murine CD4+ T cells that LMS reduced IL-6/STAT3 and TLR signaling. In agreement, we show that LMS decreased the proliferation of activated T cells, whereas we found both IL-6/STAT3 and TLR signaling gene pathways upregulated in LMS-treated T cells (Supplemental Fig 3E).

In our study, LMS induced cell cycle arrest through the induction of p53, resulting in decreased T cell activation and proliferation, without inducing cell death (Fig 3H and Supplemental Fig 1B). Induction of cell-cycle arrest - but not cell death - is in line with a modest p53 activation through suppression by negative regulators (60, 61). This arrest without cell death allows cells to engage in DNA damage repair (62). The activation of p53 can occur through multiple mechanisms, including cellular stressors such as DNA damage, hypoxia and toxic metabolites. During DNA damage, the damage sensing kinases ATM and ATR phosphorylate histone H2AX and activate downstream kinases Chk1 and Chk2 (63, 64). This intra-S-checkpoint, once activated, induces cell cycle arrest through p53. The clear mid-S-phase arrest in our cell cycle data is indicative for intra-S-checkpoint activation which is supported by the increase in pChk1 in LMS treated T cells. Furthermore, we show that a 48 hour treatment with LMS increases γ-H2AX foci, indicating the occurrence of DNA damage.

DNA damage can occur due to extracellular and intracellular stressors such as reactive oxygen species (ROS), metabolites and direct drug effects such as intercalation (41). Albendazole, an antihelminticum similar to LMS in function and structure, has been shown to be a potent DNA intercalator capable of inducing direct DNA damage (65). In our study, LMS showed relatively low intercalation abilities, and only in high dosages (Supplemental Fig 3F), making a direct intercalation mechanism in our experiments unlikely. Similarily to our data, the compound 4d, a synthetic manufactured metabolite of LMS, has been shown to induce cell-cycle arrest in HeLa cells and was found to induce DNA damage, although the direct mechanism remains unknown (66).

Increasing the understanding of the mechanism of a drug will lead to a more effective treatment. Overall, this study strengthens the idea that LMS affects human T cells directly and acts as an immunosuppressive drug. These clear immunosuppressive effects can explain its effectiveness in the treatment of INS (6). In INS, the high dosages of prednisolone needed to induce and retain remission cause serious side effects related to steroid overexposure (67). LMS has been associated with relatively mild side effects (68, 69) and could therefore be a viable adjuvant immune suppressant option in the treatment of INS. Despite these promising insights into the mechanism of LMS, questions on the exact mechanism remain. A limitation of this study is that *in vitro* experiments, although performed on human cells, cannot mimic the *in vivo* situation entirely. For this reason our project is currently studying the immunological effects of LMS treatment in INS patients. Future research will shed more light on its immunosuppressive effects on T cells and other cell types, as well as its immunological effects in patients.

## Abbreviations used

LMS: Levamisole
TNAP: Tissue non-specific alkaline phosphatase
nAChR: Nicotinic acetylcholine receptor
INS: Idiopathic nephrotic syndrome
Th2: T-helper 2 cell
Th1: T-helper 1 cell
TLR: Toll-like receptor
PBMC: Peripheral blood mononucleaur cells
MACS: Magnetic-activated cell sorting
Q-VD-Oph: Quinoline-Val-Asp-Difluorophenoxymethylketone
RIN: RNA integrity number
Log2FC: Log2 fold change
GSEA: Gene Set Enrichment Analysis
FMO: Fluorescence minus one
7-AAD: 7-aminoactinomycin-D
CFSE: CellTrace carboxyfluorescein succinimidyl ester
PFA: Paraformaldehyde
BrdU: Bromodeoxyuridine
MFI: Median fluorescent intensity
PCA: Principal component analysis
Chk1: Checkpoint kinase 1
DSB: DNA double strand breaks
5-FU: 5-fluorouracil

## 5. DATA AVAILABILITY STATEMENT

The data that is presented in this study are accessible in the Gene Expression Omnibus (GEO), accession number: GSE230683.

## 6. ETHICS STATEMENT

Buffy coats form anonymous healthy blood donors were obtained from the Sanquin Blood Supply, Amsterdam, The Netherlands. All donors provided written informed consent for usage of the remainder of their routine blood donation in research. All experiments were performed according to the ethical standards of the institutional medical ethical committee of the Amsterdam UMC, as well as in agreement with the declaration of Helsinki as revised in 1983.

## 7. AUTHOR CONTRIBUTIONS

GHK, JEJG and SF designed the study and wrote the manuscript. Experiments and analyses were performed by GHK, NC and LB. MS and GHK performed analyses on the sequencing dataset. RAH wrote the algorithm for analyzing the nuclear foci and RAH and GHK performed the foci analyses on microscopy data. FV and AHMB provided theoretical and technical input. JEJG and SF supervised the study. All authors read and approved the final manuscript.

## 9. ACKNOWLEDGEMENTS

The authors would like to thank Berend Hooibrink, Kim Brandwijk-Paarlberg and Toni van Capel for their excellent flowcytometry support. The authors would also like to thank dr. Alessandra Tammaro for providing extensive theoretical and technical input.

## 13. SUPPLEMENTAL DATA LEGENDS

<u>SUPPLEMENTAL FIGURE 1:</u> Gating strategy flow cytometry and LMS toxicity and titration. (A) FlowJo scatterplots showing the general flow cytomtery gating strategy. Gates were drawn based on fluorescence minus one (FMO) control stainings. (B) Bar graph showing the percentages of 7AAD negative events (live cells) for several LMS concentrations (n = 1 donor, replicates = 3). Activated T cells were treated for 72 hours. (C) Division Indices calculated from the CFSE cell proliferation dilution assays for different LMS concentrations are shown. Treatment with 1 mM LMS resulted in a decreased T-cell proliferation, while the lower concentrations had no effect (n = 1 donor, replicates = 3). (D) Bar graphs depicting the percentages of CD4+ (left panel) and CD8+ (right panel) T cells in solvent-treated (Ctrl) and 1 mM LMS-treated samples (n = 6 donors).

<u>SUPPLEMENTAL FIGURE 2:</u> Principal component analysis, correlation map and complete KEGG gene set enrichment analysis. (A) Principal component analysis (PCA) of the gene expression profiles of human activated T cells treated for 24 and 48 hours with 1mM LMS or solvent (control) (n = 3 donors). (B) Sample Correlation Map (SCM) depicting gene expression log2 z-scores (blue = - 1, no overlap; red = 1, maximal overlap) of the RNA-seq samples, as indicated. (C) GSEA for the KEGG pathways set, showing the gene pathways that were activated and supressed during LMS treatment. Gene pathways were considered statistically significant when FDR < 0.2 and P value <0.05.

<u>SUPPLEMENTAL FIGURE 3:</u> FasL expression, γ-H2AX expression, Gene set enrichment analysis and intercalation analysis.(A) Flowcytometry plot of FasL staining in human activated T cells treated for 72 hours with 1mM LMS (red histogram) versus solvent (black histogram) (n = 1 donor, replicates = 3). (B) Bar graph depicting the fluorescence geometric means of FasL flowcytometry stainings. Means and SDs are represented (C) Analysis of γ*-*H2AX foci in activated T cells treated with 1 mM LMS, solvent or 1 μM doxorubicin (positive control, n = 3 donors). LMS-treated samples and solvent controls were treated for 48 hours. As a positive control, cells were treated with 1 μM doxorubicin for 1 hour. (D) γ-H2AX foci in (from left to right) solvent controls, 1 mM LMS and 1 μM doxorubicin. (E) GSEA enrichment plots show significant enrichment of genes involved in IL-6/JAK/STAT3 and Toll-like receptor signaling in LMS-treated T cells. GSEA enrichment plots for (from left to right) HALLMARK_IL6_JAK_STAT3_SIGNALING and KEGG_TOLL_LIKE_RECEPTOR_SIGNALING. False discovery rate (FDR), enrichment score (ES), normalized enrichment score and P value are shown in the individual plots. (F) Bar graph depicting ethidium bromide (EtBr) fluorescence of competitive EtBr displacement assay on calf thymus DNA treated with by solvent control (Ctrl), 1 mM LMS or 10 μM doxorubicin (dox). Calf thymus DNA was pre-incubated with solvent control, 1 mM LMS or 10 μM doxorubicin for 1 hour at room temperature, after which EtBr (12.6 μM) was added and fluorescence was measured. Treatment with 10 mM LMS showed a slight reduction of EtBr fluorescence, whereas lower concentrations (1 μM – 5 mM) did not displace Etbr (**** P<.00001; One Way Analysis of variance, ANOVA with Dunnett’s multiple comparison test).

