## Supplementary figures and images for "Levamisole suppresses activation and proliferation of human T cells by the induction of a p53-dependent DNA damage response"

### Supplemental figure 1

#### Supplemental 1

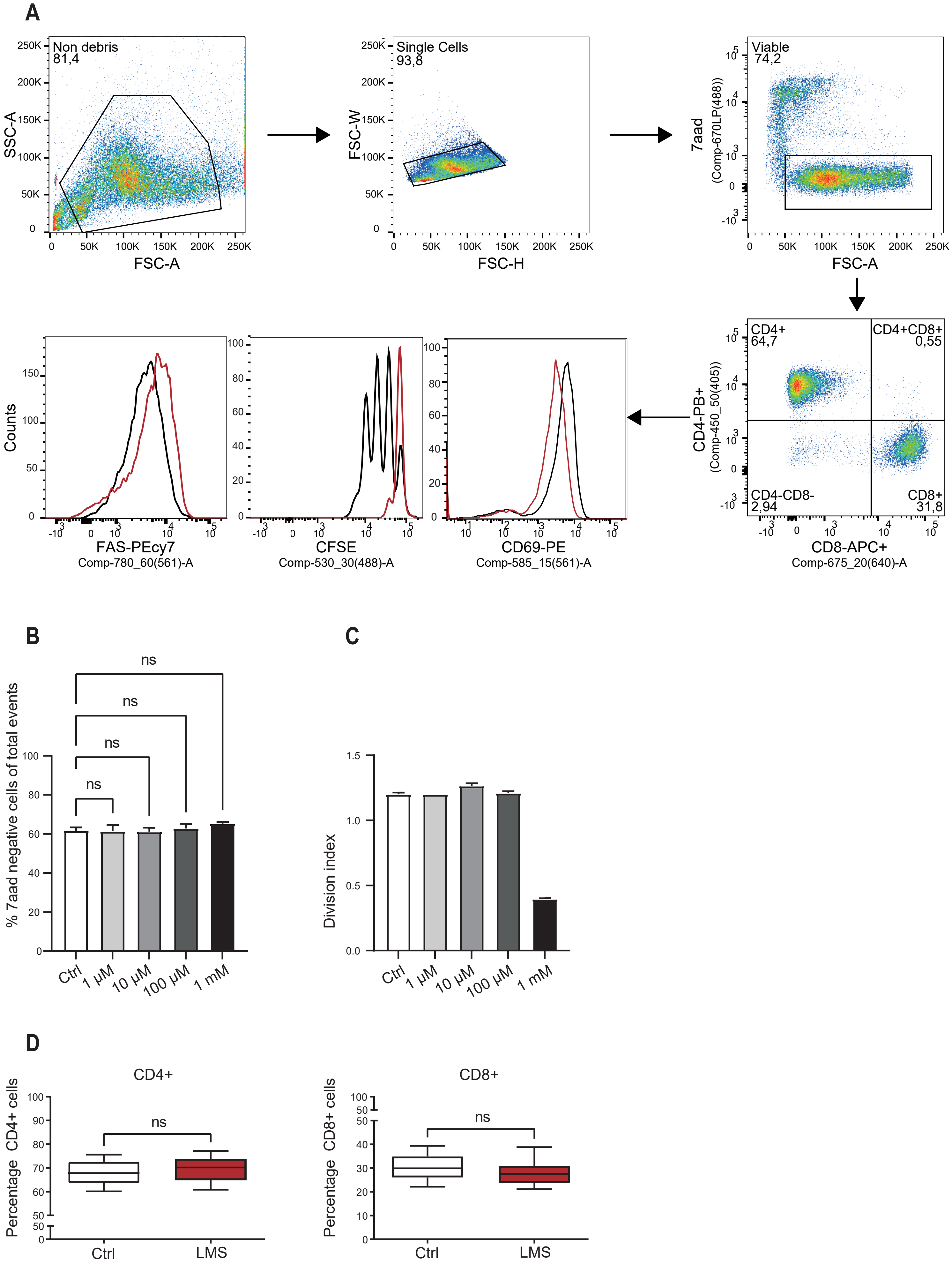

### Supplemental figure 2

#### Supplemental 2

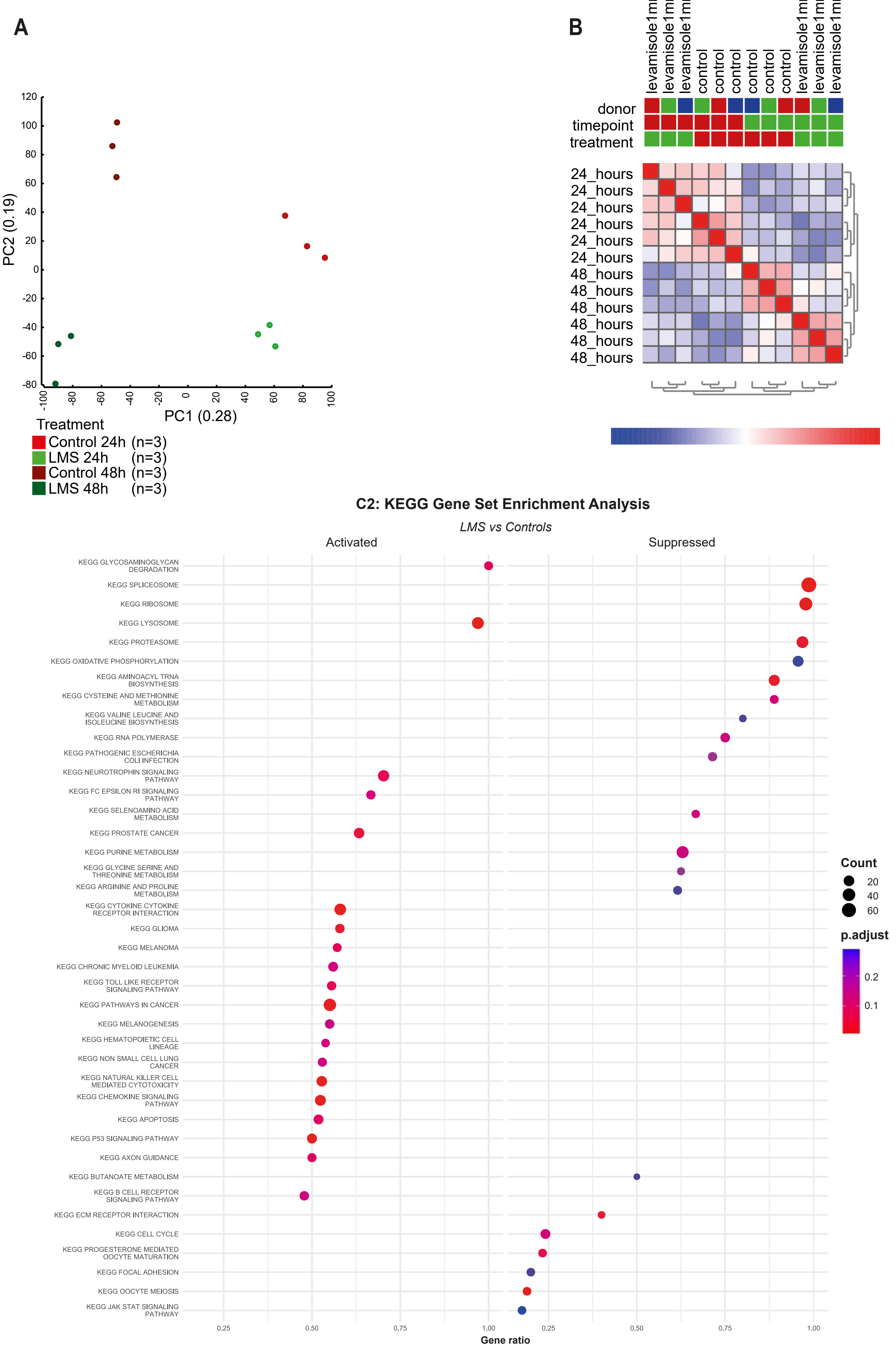

### Supplemental figure 3

#### Supplemental 3

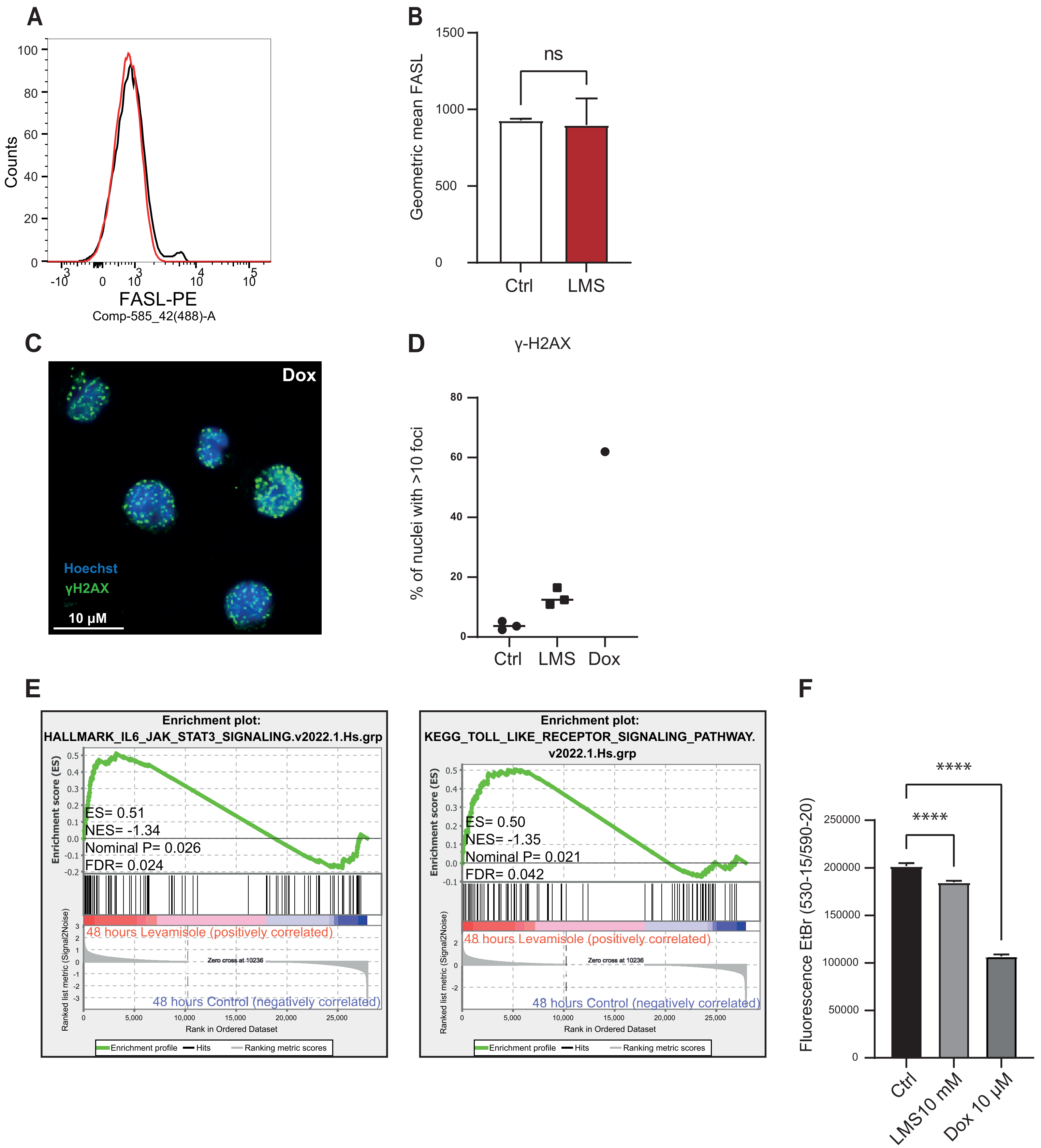
